# RNAModR: Functional analysis of mRNA modifications in R

**DOI:** 10.1101/080051

**Authors:** Maurits Evers, Andrew Shafik, Ulrike Schumann, Thomas Preiss

## Abstract

**Motivation:** Research in the emerging field of epitranscriptomics is increasingly generating comprehensive maps of chemical modifications in messenger RNAs (mRNAs). A computational framework allowing a reproducible and standardised analysis of these mRNA modification data is missing, but will be crucial for reliable functional meta-gene analyses and cross-study comparisons.

**Results:** We have developed RNAModR, an open-source and R-based set of methods, to analyse and visualise the transcriptome-wide distribution of mRNA modifications. RNAModR allows the statistical evaluation of the mRNA modification site distribution relative to null sites on a meta-gene level, providing insight into the functional role of these mRNA modifications on e.g. mRNA structure and stability.

**Availability and implementation:** RNAModR is available under the GNU General Public License (GPL) as an R-package from https://github.com/mevers/RNAModR.

## Introduction

Various chemical nucleoside modifications have been shown to occur in messenger RNAs (mRNAs). Emerging evidence suggests that these mRNA modifications may have critical regulatory roles inside cells by affecting RNA structure and the recruitment of RNA-binding proteins. This in turn may potentially affect all steps of mRNA metabolism, from splicing of pre-mRNAs to mRNA localisation, translation and stability (see e.g. [7, 6], and references therein). Two prominent examples of modifications that have been detected in mRNAs include N^6^-methyladenosine (m^6^A) and 5-methylcytidine (m^5^C). Novel experimental methods coupled with high-throughput sequencing allow transcriptome-wide profiling of various mRNA modifications, often at single-nucleotide resolution, establishing the new research field of epitranscriptomics (reviewed in e.g. [6], see also [8, 1, 3, 4]). As the list of mRNA modifications continues to increase (both in terms of numbers of identified sites and modification type), so does the need for a standardised framework allowing for a consistent and reproducible analysis of these modifications.

We introduce RNAModR, an R package that allows the reproducible statistical analysis and visualisation of RNA modifications on the transcriptome level, providing insight into the functional role of these RNA modifications on e.g. local RNA structure, mRNA stability, translation etc. A comparison with published results involving different RNA modification data demonstrates that RNAModR reproduces key findings, and therefore represents a robust analysis framework for the reproducible meta-gene analysis involving RNA modification data.

## Methods

RNAModR has been implemented as a set of tools available through a package within the statistical computing environment R ([5]), and is compatible with R versions 3.0 or later. RNAModR depends on a list of existing R/Bioconductor packages that facilitate the mapping of genomic to transcriptomic loci and provide methods for handling genome and transcriptome annotations and sequences. Specific details are provided in the RNAModR package manual, in the supplementary materials and on the public github project page. The package can be downloaded and installed directly from https://github.com/mevers/RNAModR. As minimal input, RNAModR requires a list of genomic loci of RNA modifications in the BED (browser extensible data) file format.

Within RNAModR we provide sample data compiled from transcriptome-wide profiling studies of different RNA modifications in human cell lines: N(1)-methyladenosine (m^1^A) modifications from [2], N^6^-methyladenosine (m^6^A) and N^6^,2’-*O*-dimethyladenosine (m^6^Am) modifications from [4], and 5-methylcytidine (m^5^C) modifications from [8]. All sample data are included as BED files, with genome coordinates based on the GRCh38/hg38 genome assembly version; we provide R scripts in the supplementary materials to reproduce results shown here. We emphasise that RNAModR is not limited to methylation-type modification data, but can be applied to other RNA modification data such as pseudouridine, A-to-I editing etc.

We discuss key steps in a typical RNAModR workflow analysing a set of RNA modifications provided as a BED file of genomic loci.

### Transcriptome construction

In a first step RNAModR generates a custom reference transcriptome (function BuildTx()). The function automatically downloads a UCSC RefSeq-based reference genome and annotation, and constructs a custom transcriptome. To allow for unique identification of sites with transcripts and to avoid double-counting of sites located in multiple transcript isoforms, we collapse the transcriptome by choosing for every gene the transcript isoform with the longest CDS (coding sequence) and longest adjoining 5’/3’UTRs (untranslated regions) among the set of transcript isoforms. The resulting canonical transcriptome is stored locally for further downstream analysis, and contains position and sequence information on non-overlapping transcript sections per gene; “transcript sections” are defined as a promoter region (up to 1000 nt upstream of the 5’UTR start site), 5’UTR, CDS, 3’UTR, and introns. To guarantee reproducibility of results, the same reference transcriptome can and should be used for additional analyses. RNAModR currently supports human and mouse genomes based on the two latest genome assembly versions (human: hg38, hg19; mouse: mm10, mm9), but can be extended to allow for other organisms.

### Mapping to transcriptome

We use the canonical transcriptome to map genomic loci of RNA modifications to positions within different transcript sections (function `SmartMap(…)`). We provide visualisation routines to assess the distribution of sites across and within different transcript sections (functions `PlotSectionDistribution(…), PlotSpatialDistribution(…))`.

### Functional analyses

Functional analyses aim at providing insight into spatial or functional associations of RNA modifications with particular transcript features (e.g. transcript sections, GC content, splice-sites), and/or secondary data (e.g. RNA-binding protein target sites etc.). For example, an enrichment analysis may explore the enrichment or depletion of RNA modifications within a particular transcript section, or whether RNA modifications localise with particular transcript features such as the start codon, stop codon, etc. In order to do so, we have implemented statistical methods (based on multiple Fisher’s exact tests) to assess the enrichment of observed sites relative to a list of sites based on a suitable null distribution. The latter can be provided manually as a list of empirical “null” sites based on alignment data (for example a list of all *non*-modified nucleotides of the same type), or can be generated automatically using two different approaches: In a nucleotide abundance-specific approach (function `GenerateNull(…, method = “ntAbund”, nt = …)` we use the position information and abundance of all non-modified nucleotides of type *N* in transcript sections that contain at least one modified nucleotide mod*N*; a permutation-based approach (function `GenerateNull(…, method = “perm”)`) generates a set of null sites by uniform-randomly permuting the position of a modified nucleotide within the corresponding transcript section. It is important to note that the choice of null sites might critically affect the results of the enrichment analyses; in the case of large non-uniformity of nucleotide abundances within transcript sections, we strongly recommend using alignment data-derived null sites or null sites inferred from the nucleotide abundance-specific approach. Please see also the supplementary materials for further details involving the statistical enrichment analysis.

Spatial enrichment across and within transcript section can be assessed and visualised (functions `PlotSectionEnrichment(…), PlotSpatialEnrichment(…)`; furthermore, RNAModR provides routines to assess and visualise the spatial enrichment of sites relative to other transcript-level features such as e.g. exon-exon junctions or secondary user-provided data (functions `PlotRelDistDistribution(…), PlotRelDistEnrichment(…))`. RNAModR already includes sample data of key RNA-binding protein footprints and miRNA target sites, relevant to the analysis of the included RNA modification data. Details on the sample data are given in the supplementary materials.

We also provide routines to visualise nucleotide abundances within a user-defined window centred around the RNA modification and “null” sites. RNAModR can visualise corresponding sequence logos (function `PlotSeqLogo(…)`), and evaluate and visualise the difference in the distribution of GC content using a two-tailed two-sample t-test (function `PlotGC(…)`.

## Results and conclusion

We compare RNAModR results with published results involving m^1^A, m^6^A/m^6^Am and m^5^C modification data in Figures 1–6 in the supplementary materials. RNAModR independently and successfully confirms results presented in [4, 2, 8], and provides an easy-to-use R-based package to (1) perform reproducible transcriptome-level functional analyses of RNA modifications, and (2) produce publication-ready figures.

**Figure 1:**
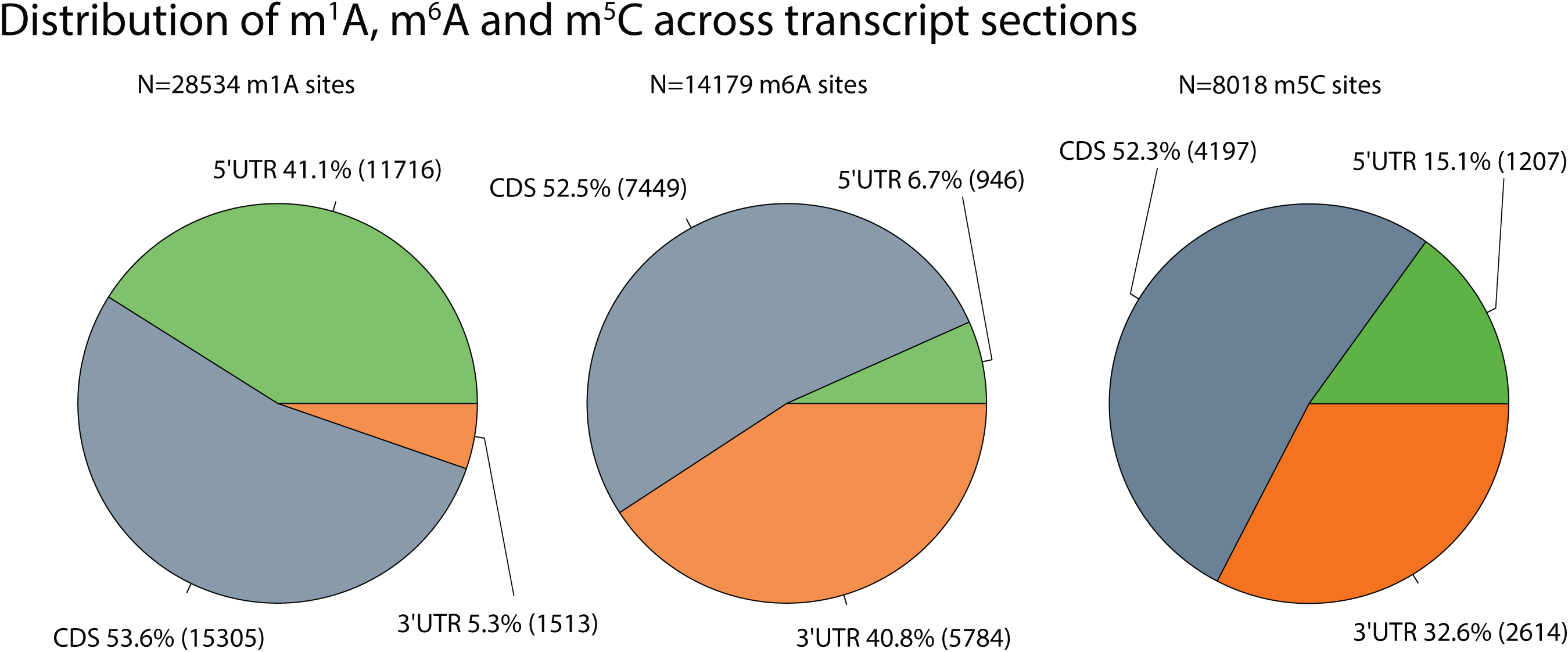
Distribution of RNA modifications across different transcript sections. Results show excellent agreement with results shown in the original publications.

**Figure 2:**
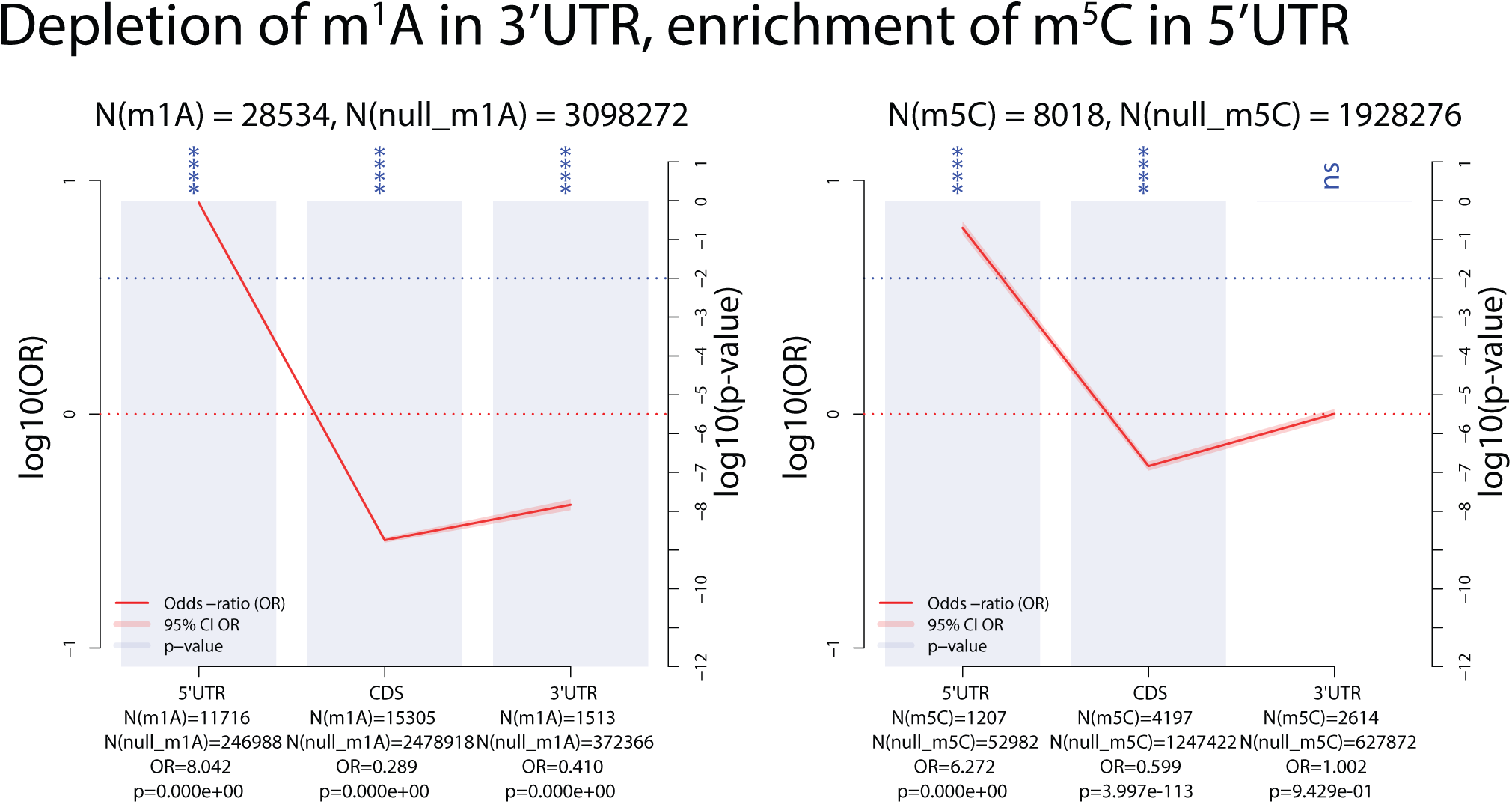
Results confirming the depletion of m^1^A in the 3’UTR, and enrichment of m^5^C in the 5’UTR (see [3, 8]).

**Figure 3:**
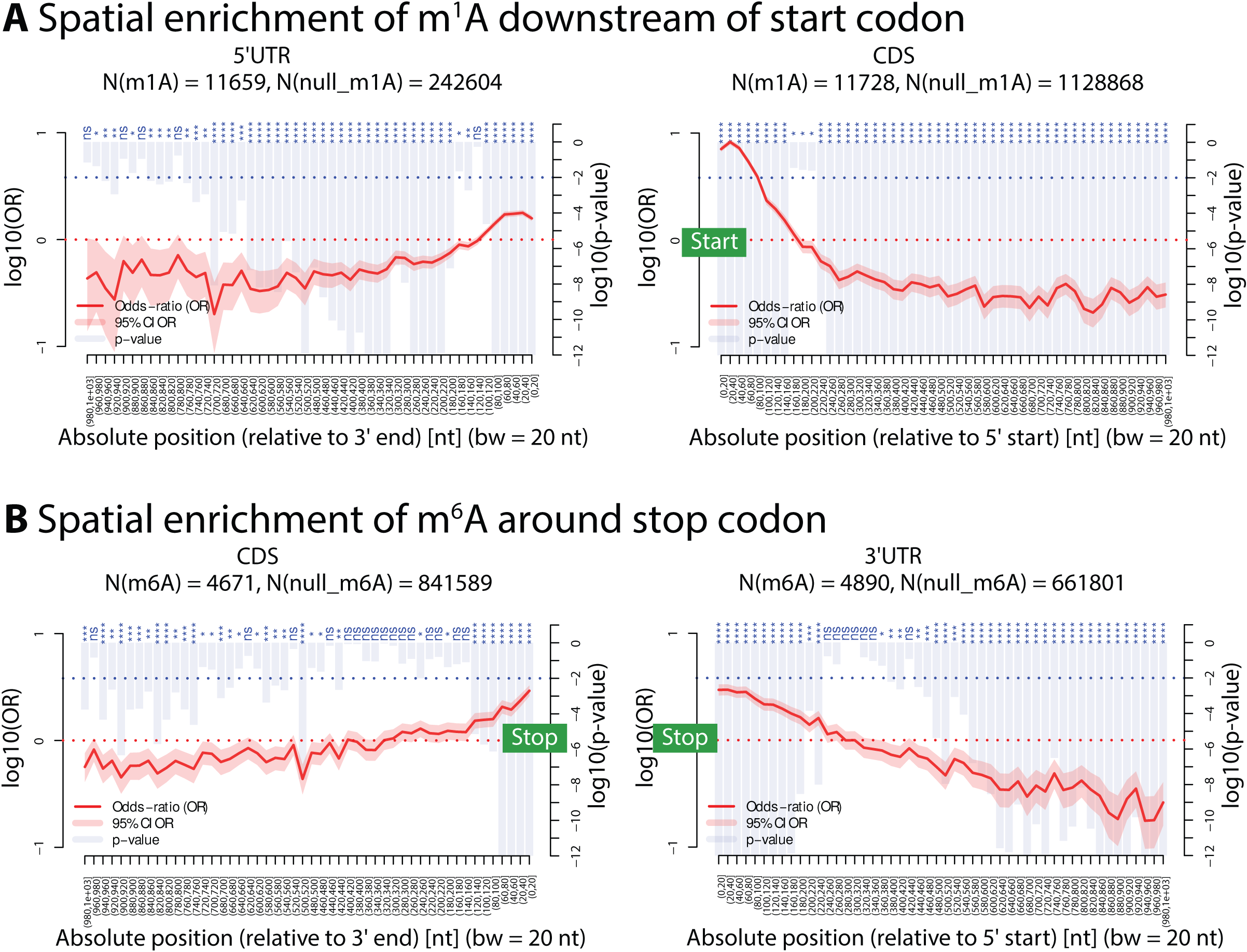
**A**: Results confirming the spatial enrichment of m^1^A within the first 100 nt downstream of the start codon in the CDS (see [3]). **B**: Results confirming the enrichment of m^6^A around the stop codon (upstream into the CDS and downstream into the 3’UTR (see [3]). The location of start and stop codon are labelled by the green start and stop textboxes.

**Figure 4:**
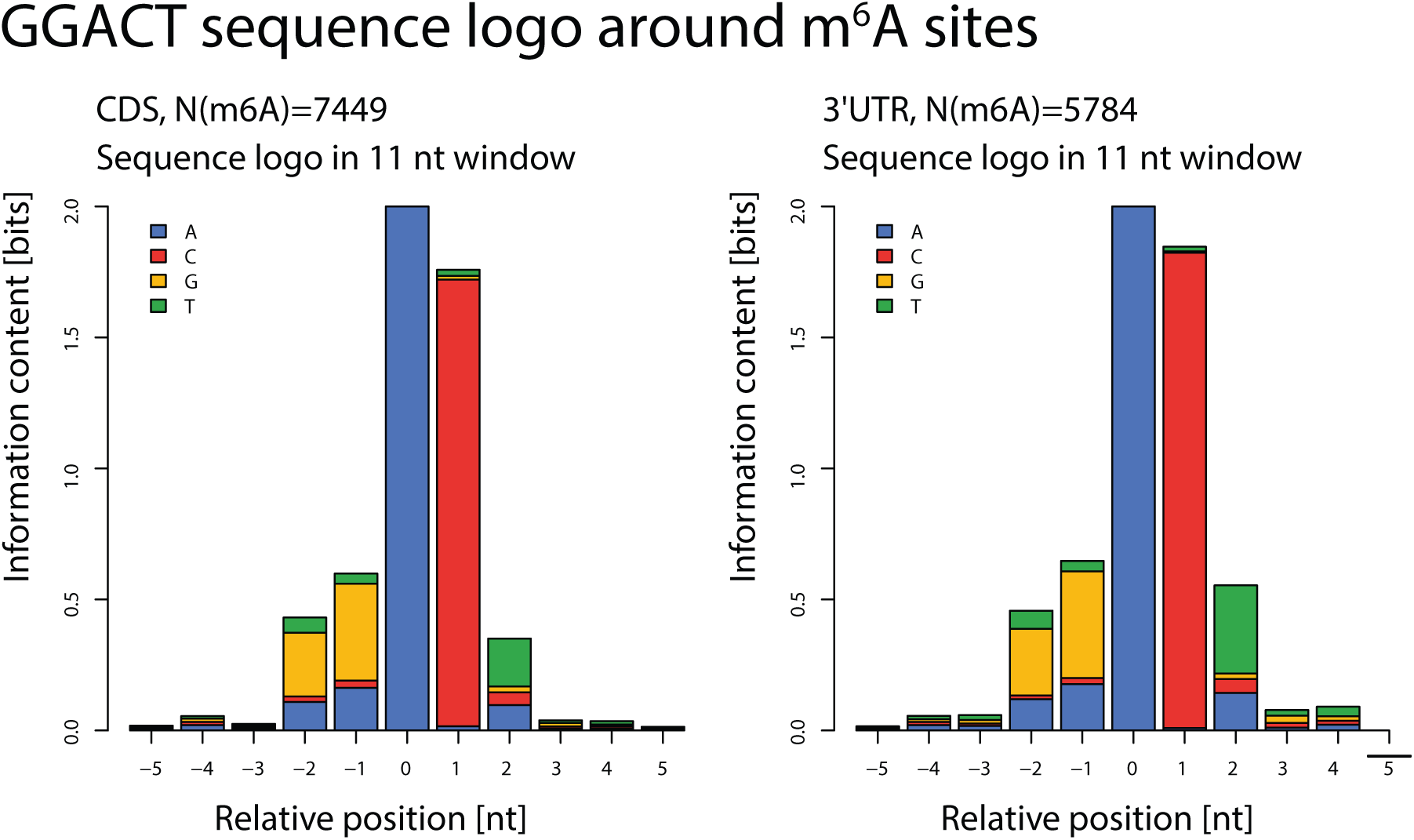
Nucleotide abundances within a 21 nt window around m^6^A sites in the CDS and 3’UTR, confirming the GGACT motif (see [3]).

**Figure 5:**
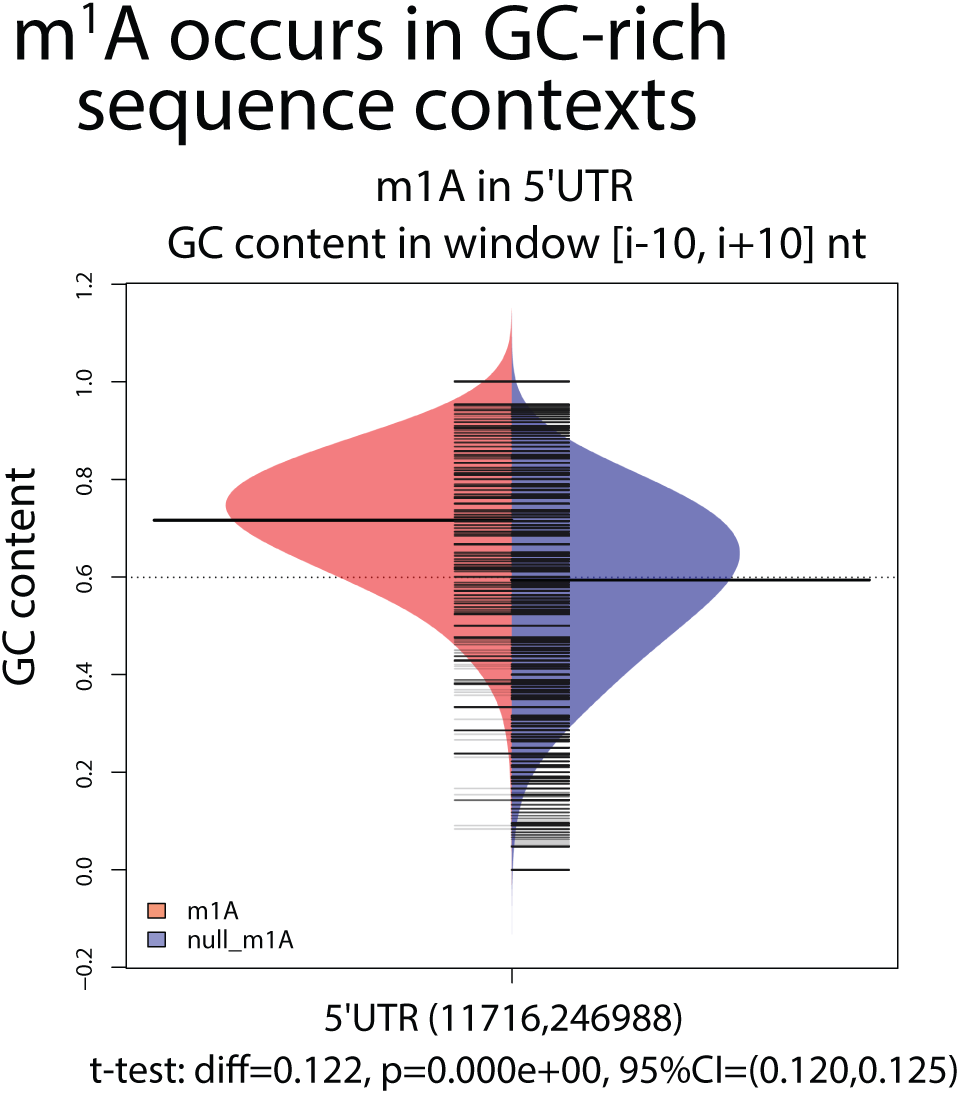
RNAModR confirmation that m1A occurs in GC-rich regions in the 5’UTR (see [3]).

**Figure 6:**
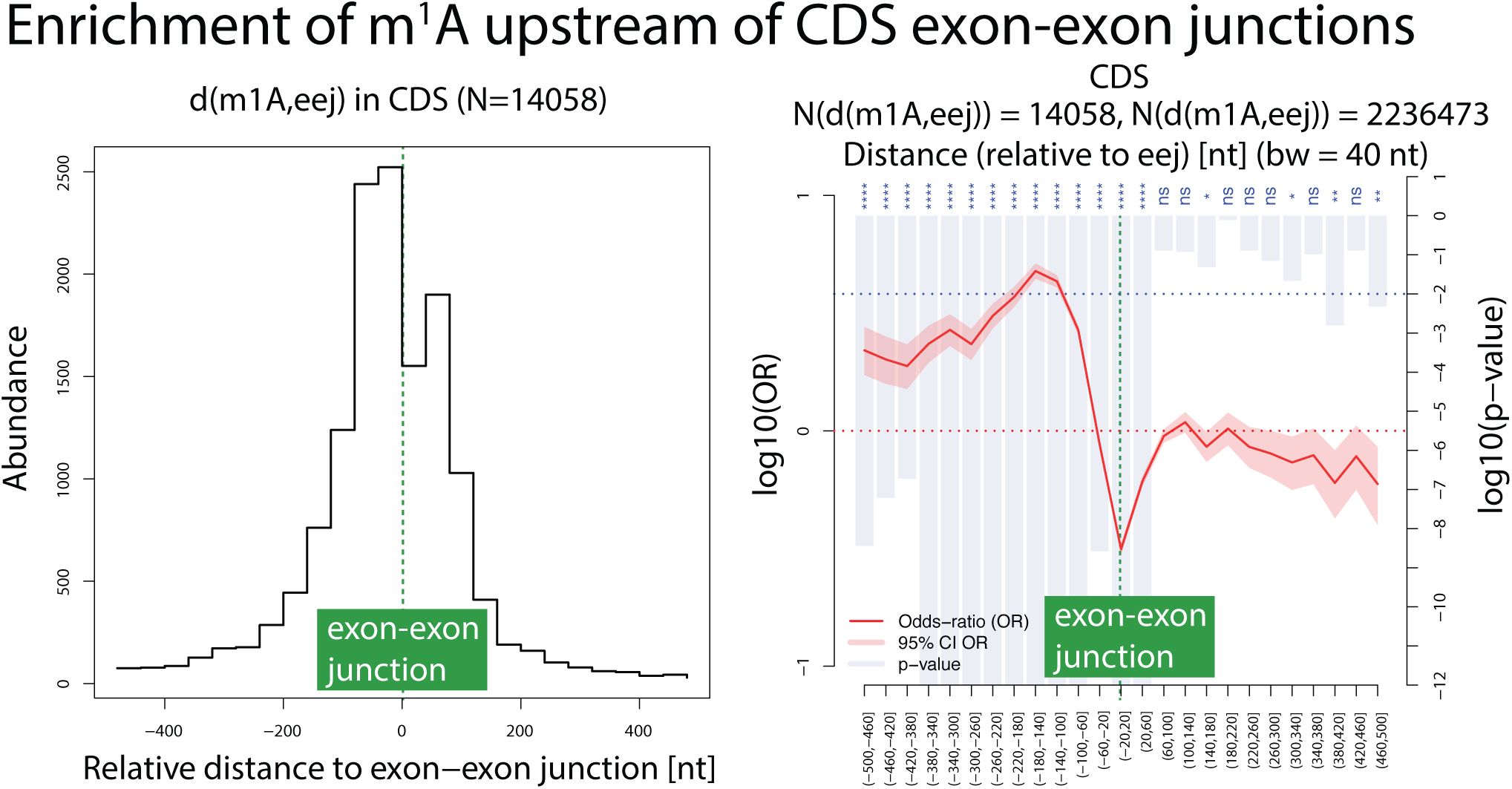
Results confirming an enrichment of m^1^A sites upstream of the nearest exon-exon junction in the CDS (see [3]). The reference position of the exon-exon junctions is marked by the green dashed line and labelled as indicated.

## Supplementary materials for RNAModR: Functional analysis of mRNA modifications in R

### 1 Summary plots for the RNAModR-based analysis of mRNA modification data

Data are based on the following original studies:

1. m^1^A from Ref. [3]
2. m^6^A/m^6^Am from Ref. [5]
3. m^5^C from Ref. [8]

All plots were generated using RNAModR functions, with minimal post-processing to increase readability. Section 2 provides additional details involving the quantities shown in Figures 1–6.

### 2 Details on key quantities shown in abundance and enrichment plots

#### 2.1 Abundance plots

An example of a typical spatial abundance plot is shown in the following figure, based on m^6^A data from [5]:

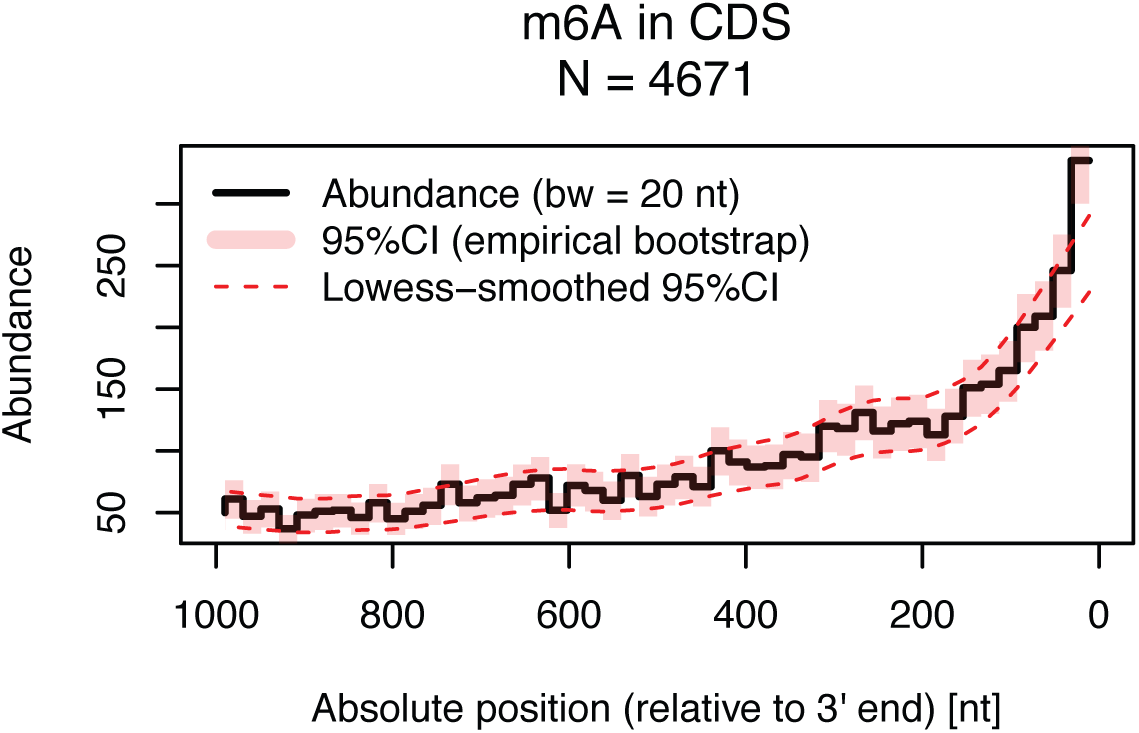

We summarise key aspects and quantities of the plot:

- Site abundances are binned and shown as the number of sites per bin across a fixed window. Both the window size and bin width can be changed by the user, see the corresponding help files in R (e.g. `?PlotSpatialDistribution`). In above figure, the window size is 1000 nt, and the bin width 20 nt.
- The number in the title (in above figure, `N = 4671`) refers to the total number of sites within the shown window.
- By default, spatial abundance plots are shown with 95% confidence intervals (CI). CI’s are derived from an empirical bootstrap procedure by sampling with replacement from the spatial data. The default sampling size is 5000 and can be changed by the user. Corresponding empirical bootstrap-based 95% CI’s are shown as red shaded areas surrounding the abundance values per bin; the dashed red lines show Lowess-smoothed 95% CI’s.

#### 2.2 Enrichment plots

An example of a typical spatial enrichment plot is shown in the following figure, based on m^6^A data from [5]. Null sites correspond to all non-methylated adenosines from transcripts that contain at least one m^6^A site (generated using the RNAModR function negSites <- `GenerateNull(posSites, method = “ntAbund”, nt = “A”)`, where posSites are the transcriptome-mapped m^6^A sites):

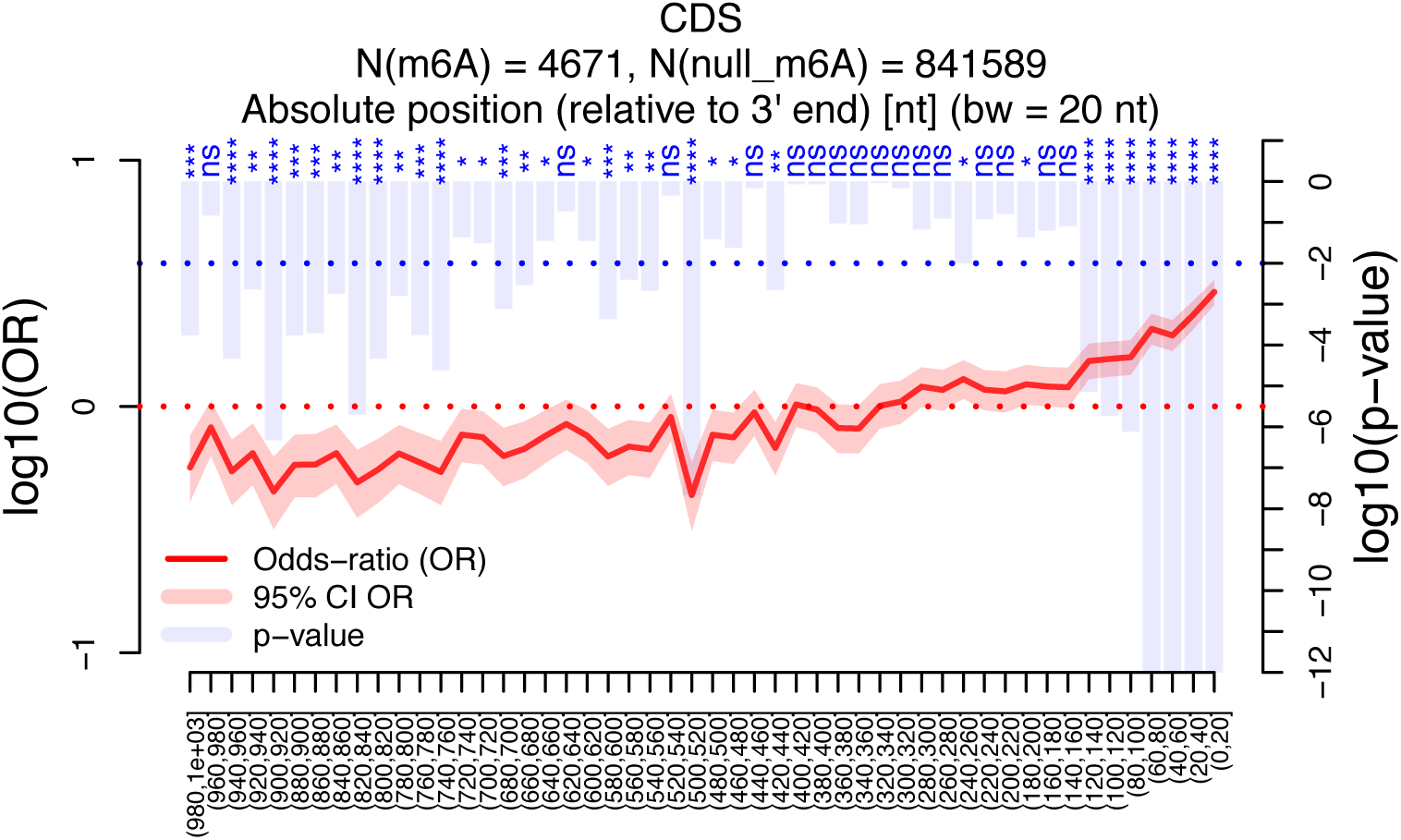

We summarise key aspects and quantities of the plot:

- The solid red line shows log10-transformed odds-ratios (OR’s), with the corresponding y-axis shown on the left side of the plot. The OR in bin *i* is calculated as

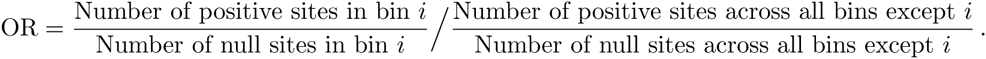 Shaded red areas indicate 95% confidence intervals of the OR’s, obtained from assessing the statistical significance of OR using Fisher’s exact test.
- Statistical significance is assessed using Fisher’s exact tests against a null hypothesis of OR = 1. Resulting multiple Fisher’s exact tests give a *p*-value for every OR per bin. We correct raw *p*-values for multiple hypothesis testing using the method of Benjamini and Hochberg [2]. Resulting multiple-testing adjusted and log10-transformed *p*-values are shown by the blue bars, with the corresponding y-axis shown on the right side of the plot. *p*-values are shown in the range [10^−12^ … 1]. Additionally, statistical significance using the “star-notation” is indicated above every bar:

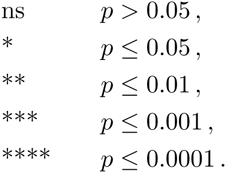
- Values on the x-axis give the ranges per bin, with a round (square) bracket indicating an exclusive (inclusive) value. Equivalently to abundance plots, both the window size and bin width can be changed by the user, see the corresponding help files in R (e.g. `?PlotSpatialEnrichment`). In above figure, the window size is 1000 nt, and the bin width 20 nt.
- The numbers in the title give the the total number of positive sites (in above figure, `N(m6A) = 4671`) and null sites (in above figure, `N(null m6A) = 841589`) within the shown window. The last line in the title may also contain details on the spatial reference point (if applicable), e.g. in above figure, spatial positions are relative to the 3’ end of the CDS.

### 3 Sample data included in RNAModR

All sample data are provided in the BED (browser extensible data) file format. Genomic coordinates are based on the human hg38 reference genome.

1. List of m^1^A modifications from Ref. [3]
2. List of m^6^A modifications from Ref. [5]
3. List of m^5^C modifications from Ref. [8]
4. List of eIF3 target loci from Ref. [6]
5. List of eIF4AIII target loci from Ref. [7]
6. List of YTHDF2 target loci from Ref. [9]
7. List of AGO1-4 target loci from Ref. [4]
8. List of TargetScan-based miRNA target loci [1]

### 4 R script to reproduce all figures

~~~
# Load the RNAModR library
library(RNAModR);
# Build transcriptome
BuildTx("hg38");
###########################################################
#################### m6A data analysis ####################
###########################################################
# Read in m6A data from Linder et al.
bedFile <- system.file("extdata",
"miCLIP_m6A_Linder2015_hg38.bed",
package = "RNAModR");
sites <- ReadBED(bedFile);
# Map data to transcriptome
posSites <- SmartMap(sites, id = "m6A", refGenome = "hg38");
# Generate null sites
negSites <- GenerateNull(posSites, method = "ntAbund", nt = "A");
# Filter sites that overlap with 5’UTR, CDS, 3’UTR
posSites <- FilterTxLoc(posSites, filter = c("5’UTR", "CDS", "3’UTR"));
negSites <- FilterTxLoc(negSites, filter = c("5’UTR", "CDS", "3’UTR"));
# Distribution of sites across transcript sections
PlotSectionDistribution(posSites);
# Spatial distribution of sites across transcript sections
PlotSpatialDistribution(posSites, nbreaks = 50, absolute = TRUE);
# Spatial enrichment of sites across transcript sections
PlotSpatialEnrichment(posSites, negSites, binWidth = 20, posMax = 1000);
# Sequence logo within 11 nt window around m6A sites
PlotSeqLogo(posSites);
###########################################################
#################### m1A data analysis ####################
###########################################################
# Read in m1A data from Dominissini et al.
bedFile <- system.file("extdata",
          "MeRIPseq_m1A_Dominissini2016_hg38.bed",
          package = "RNAModR");
sites <- ReadBED(bedFile);
# Map data to transcriptome
posSites <- SmartMap(sites, id = "m1A", refGenome = "hg38");
# Generate null sites
negSites <- GenerateNull(posSites, method = "ntAbund", nt = "A");
# Filter sites that overlap with 5’UTR, CDS, 3’UTR
posSites <- FilterTxLoc(posSites, filter = c("5’UTR", "CDS", "3’UTR"));
negSites <- FilterTxLoc(negSites, filter = c("5’UTR", "CDS", "3’UTR"));
# Distribution of sites across transcript sections
PlotSectionDistribution(posSites);
# Spatial distribution of sites across transcript sections
PlotSpatialDistribution(posSites, nbreaks = 50, absolute = TRUE);
# Spatial enrichment of sites across transcript sections
PlotSpatialEnrichment(posSites, negSites, binWidth = 20, posMax = 1000);
# GC content analysis within a 21 nt window
PlotGC(posSites, negSites, flank = 10);
###########################################################
#################### m5C data analysis ####################
###########################################################
# Read in m1A data from Squires et al.
bedFile <- system.file("extdata",
"bsRNAseq_m5C_Squires2012_hg38.bed",
package = "RNAModR");
sites <- ReadBED(bedFile);
# Map data to transcriptome
posSites <- SmartMap(sites, id = "m5C", refGenome = "hg38");
# Generate null sites
negSites <- GenerateNull(posSites, method = "ntAbund", nt = "C");
# Filter sites that overlap with 5’UTR, CDS, 3’UTR
posSites <- FilterTxLoc(posSites, filter = c("5’UTR", "CDS", "3’UTR"));
negSites <- FilterTxLoc(negSites, filter = c("5’UTR", "CDS", "3’UTR"));
# Distribution of sites across transcript sections
PlotSectionDistribution(posSites);
# Spatial distribution of sites across transcript sections
PlotSpatialDistribution(posSites, nbreaks = 50, absolute = TRUE);
# Spatial enrichment of sites across transcript sections
PlotSpatialEnrichment(posSites, negSites, binWidth = 20, posMax = 1000);
~~~

## References

[1] D. Dominissini, S. Moshitch-Moshkovitz, S. Schwartz, M. Salmon-Divon, L. Ungar, S. Osenberg, K. Cesarkas, J. Jacob-Hirsch, N. Amariglio, M. Kupiec, R. Sorek, and G. Rechavi. Topology of the human and mouse m6A RNA methylomes revealed by m6A-seq. Nature, 485(7397):201–206, May 2012.

[2] D. Dominissini, S. Nachtergaele, S. Moshitch-Moshkovitz, E. Peer, N. Kol, M. S. Ben-Haim, Q. Dai, A. Di Segni, M. Salmon-Divon, W. C. Clark, G. Zheng, T. Pan, O. Solomon, E. Eyal, V. Hershkovitz, D. Han, L. C. Doré, N. Amariglio, G. Rechavi, and C. He. The dynamic N(1)-methyladenosine methylome in eukaryotic messenger RNA. Nature, 530(7591):441–446, Feb. 2016.

[3] S. Hussain, J. Aleksic, S. Blanco, S. Dietmann, and M. Frye. Characterizing 5-methylcytosine in the mammalian epitranscriptome. Genome Biology, 14(11):215, 2013.

[4] B. Linder, A. V. Grozhik, A. O. Olarerin-George, C. Meydan, C. E. Mason, and S. R. Jaffrey. Single-nucleotide-resolution mapping of m6A and m6Am throughout the transcriptome. Nature Methods, 12(8):767–772, Aug. 2015.

[5] R Core Team. R: A Language and Environment for Statistical Computing. R Foundation for Statistical Computing, Vienna, Austria, 2016.

[6] S. Schwartz. Cracking the epitranscriptome. RNA, 22(2):169–174, Feb. 2016.

[7] A. Shafik, U. Schumann, M. Evers, T. Sibbritt, and T. Preiss. The emerging epitranscriptomics of long noncoding RNAs. Biochimica et biophysica acta, 1859(1):59–70, Jan. 2016.

[8] J. E. Squires, H. R. Patel, M. Nousch, T. Sibbritt, D. T. Humphreys, B. J. Parker, C. M. Suter, and T. Preiss. Widespread occurrence of 5-methylcytosine in human coding and non-coding RNA. Nucleic Acids Research, 40(11):5023–5033, June 2012.

## References

[1] V. Agarwal, G. W. Bell, J.-W. Nam, and D. P. Bartel. Predicting effective microRNA target sites in mammalian mRNAs. eLife, 4, 2015.

[2] Y. Benjamini and Y. Hochberg. Controlling the False Discovery Rate: A Practical and Powerful Approach to Multiple Testing. Journal of the Royal Statistical Society, Series B (Methodological), 57(1):289–300, Aug. 1995.

[3] D. Dominissini, S. Nachtergaele, S. Moshitch-Moshkovitz, E. Peer, N. Kol, M. S. Ben-Haim, Q. Dai, A. Di Segni, M. Salmon-Divon, W. C. Clark, G. Zheng, T. Pan, O. Solomon, E. Eyal, V. Hershkovitz, D. Han, L. C. Doré, N. Amariglio, G. Rechavi, and C. He. The dynamic N(1)-methyladenosine methylome in eukaryotic messenger RNA. Nature, 530(7591):441–446, Feb. 2016.

[4] M. Hafner, M. Landthaler, L. Burger, M. Khorshid, J. Hausser, P. Berninger, A. Rothballer, M. Ascano, A.-C. Jungkamp, M. Munschauer, A. Ulrich, G. S. Wardle, S. Dewell, M. Zavolan, and T. Tuschl. Transcriptome-wide identi?cation of RNA-binding protein and microRNA target sites by PAR-CLIP. Cell, 141(1):129–141, Apr. 2010.

[5] B. Linder, A. V. Grozhik, A. O. Olarerin-George, C. Meydan, C. E. Mason, and S. R. Jaffrey. Single-nucleotide-resolution mapping of m6A and m6Am throughout the transcriptome. Nature Methods, 12(8):767–772, Aug. 2015.

[6] K. D. Meyer, D. P. Patil, J. Zhou, A. Zinoviev, M. A. Skabkin, O. Elemento, T. V. Pestova, S.-B. Qian, and S. R. Jaffrey. 5' UTR m(6)A Promotes Cap-Independent Translation. Cell, 163(4):999–1010, Nov. 2015.

[7] J. Saulière, V. Murigneux, Z. Wang, E. Marquenet, I. Barbosa, O. Le Tonquèze, Y. Audic, L. Paillard, H. Roest Crollius, and H. Le Hir. CLIP-seq of eIF4AIII reveals transcriptome-wide mapping of the human exon junction complex. Nature Publishing Group, 19(11):1124–1131, Nov. 2012.

[9] X. Wang, Z. Lu, A. Gomez, G. C. Hon, Y. Yue, D. Han, Y. Fu, M. Parisien, Q. Dai, G. Jia, B. Ren, T. Pan, and C. He. N6-methyladenosine-dependent regulation of messenger RNA stability. Nature, 505(7481):117–120, Jan. 2014.

